# Environmentally-induced epigenetic conversion of a piRNA cluster

**DOI:** 10.1101/360743

**Authors:** Karine Casier, Valérie Delmarre, Nathalie Gueguen, Catherine Hermant, Elise Viodé, Chantal Vaury, Stéphane Ronsseray, Emilie Brasset, Laure Teysset, Antoine Boivin

## Abstract

Transposable element (TE) activity is repressed in animal gonads by PIWI-interacting RNAs (piRNAs), a class of small RNAs produced by specific loci made of TEs insertions and fragments. Current models propose that these loci are functionally defined by the maternal inheritance of piRNAs produced during the previous generation, raising the question of their first activation in the absence of piRNAs. Taking advantage of an inactive cluster of *P*-element derived transgene insertions, we show here that raising flies at high temperature (29°C) instead of 25°C results in a rare but invasive epigenetic conversion of this locus into an active piRNAs producing one. The newly acquired epigenetic state is stable over many generations even when flies are switch back to 25°C. The silencing capacities, piRNA production and chromatin modifications of the cluster are all identical whether conversion occurred by maternal piRNA inheritance or by high temperature. We also demonstrate that in addition to high temperature, a single homologous transgene inserted elsewhere in the genome is required to activate the locus. We thus have identified a minimal system of three components to create a stable piRNA producing locus: 1) a locus with multiple TE derived sequences; 2) an euchromatic copy of these sequences and 3) elevated temperature. Altogether, these data report the first case of the establishment of an active piRNA cluster by environmental changes. It highlights how such variations of species natural habitat can become heritable and shape their epigenome.

**SIGNIFICANCE STATEMENT:** Recently, we have witnessed great progress in our understanding of the silencing of Transposable Elements (TEs) by piRNAs, a class of small RNAs produced by piRNA clusters. At each generation, piRNA clusters are supposed to be activated by homologous piRNAs inherited from the mother raising the question of the making of the first piRNAs. Here, we report the birth of a stable and functional piRNA cluster induced by high temperature without maternal inheritance of homologous piRNAs. We propose a minimal system to create a piRNA cluster: a sufficient number of repeated sequences, a euchromatic copy of these sequences and an increase in the production of antisense RNA.

## INTRODUCTION

Transposable element (TE) activity needs to be repressed to avoid severe genome instability and gametogenesis defects. In humans, growing evidences have implicated TE in several disorders such as cancers defining a new field of diseases called transposopathies (1, 2). In the animal germline, TE activity is controlled at both transcriptional and post-transcriptional levels by small RNAs called piRNAs associated with the PIWI clade of germline Argonaute proteins (Piwi, Aub and Ago3 in *Drosophila*) (3–6). piRNAs are processed from transcripts produced from specific heterochromatic loci enriched in TE fragments, called piRNA clusters (3, 4). These loci undergo non-canonical transcription, ignoring splicing and transcription termination signals licensed by specific protein complexes such as Rhino-Deadlock-Cutoff (7, 8) and Moonshiner-TRF2 (9). Thus, when a new TE inserts into a naive genome, it will freely transpose until one copy gets inserted into a piRNA cluster leading to the production of homologous new TE piRNAs that will then repress transposition (10). In support of this idea, exogenous sequences inserted into preexisting piRNA clusters lead to the production of matching piRNAs (11–15). The specificity of the efficient repression mediated by piRNAs appears to be determined solely by the piRNA cluster sequences. It raises thus the question of how piRNA cluster loci are themselves specified. Histone H3 lysine 9 tri-methylation (H3K9me3) that is recognized by Rhino, a paralog of heterochromatin protein HP1 (16), is a shared feature of piRNA clusters. However, enrichment of H3K9me3 is not specific to piRNA clusters and tethering Rhino onto a transgene leads to the production of piRNAs only when both sense and antisense transcripts are produced (8). This suggests that neither H3K9me3 marks nor having Rhino-bound is sufficient to induce piRNA production. One current model proposes that piRNAs clusters are defined and activated at each generation by the deposition in the egg of their corresponding piRNAs from the mother (17). In support of this model, we described previously the first case of a stable transgenerational epigenetic conversion known as paramutation in animals (11). This phenomenon was first described in plants and defined as “*an epigenetic interaction between two alleles of a locus, through which one allele induces a heritable modification of the other allele without modifying the DNA sequence”* (18, 19). In our previous study, we showed that an inactive non-producing piRNA cluster of *P* transgene insertions inherited from the father can be converted into a piRNA-producing cluster by piRNAs inherited from the mother (11). However, this attractive model does not answer the question of how the first piRNAs were produced.

To address this paradox, we used the same *BX2* cluster of seven *P(lacW)* transgenes which resulted from multiple and successive *P(lacW)* transposition events, thus resembling the structure of natural piRNA clusters (20). The key advantage of the *BX2* locus is that it can exist in two epigenetic states for the production of germline piRNAs: 1) the inactive state (*BX2^OFF^*) does not produce any piRNAs and thus is unable to repress the expression of homologous sequences, and 2) the active state (*BX2^ON^* also called *BX2\**) produces abundant piRNAs that functionally repress a homologous reporter transgene in the female germline (11, 12). We therefore used *BX2* in an inactive state to search for conditions that would convert it into an active piRNA-producing locus, without pre-existing maternal piRNAs. In this report, we describe how developing flies at high temperature – 29°C instead of 25°C – induces the conversion of an inactive *BX2* locus (*BX2^OFF^*) into a stable piRNA cluster exhibiting repression properties (*BX2^ON^*). It should be noted that flies in their natural habitat exhibit this range of temperature, especially in the context of global warming. These data provide the first report of a *de novo* piRNA cluster establishment independently from maternal inheritance of homologous piRNAs and underline how environmental changes can stably induced transgenerational modification of the epigenome.

## MATERIALS AND METHODS

### Transgenes and strains

The *BX2* line carries seven *P-lacZ-white* transgenes, (*P(lacW)*, FBtp0000204) inserted in tandem and in the same orientation at cytological site 50C (20). *P(TARGET)* corresponds to *P(PZ)* (FBtp0000210), a *P-lacZ-rosy* enhancer-trap transgene inserted into the *eEF5* gene at 60B7 and expressing ß-Galactosidase in the germline (FBst0011039). References of other transgenes and strains used in this study are detailed in Supplementary Information.

### ß-Galactosidase staining

Ovarian *lacZ* expression assays were carried out using X-gal overnight staining at 37°C as previously described (21). After mounting in glycerol/ethanol (50/50), the germline *lacZ* repression was then calculated by dividing the number of repressed egg chambers by the total number of egg chambers. Most of the time, the total number of egg chambers was estimated by multiplying the number of mounted ovaries by 60, corresponding to an average of 3 to 4 egg chambers per ovariole and 16 to 18 ovarioles per ovary. Pictures were acquired with an Axio-ApoTome (Zeiss) and ZEN2 software.

### Fly dissection and RNA extraction

For each genotype tested, 20 pairs of ovaries were manually dissected in PBS 1X. For small RNA sequencing, total RNA was extracted using TRIzol (Life Technologies) as described in the reagent manual (http://tools.lifetechnologies.com/content/sfs/manuals/trizol_reagent.pdf). For the RNA precipitation step, 100% ethanol was used instead of isopropanol. For RT-qPCR experiments, total RNA was extracted using TRIzol for *BX2* and *w^1118^* females or RNeasy kit (Qiagen) for *P(TARGET)* females. Up to 6 biological replicates were used for each genotype.

### small RNA sequencing analyses

Analyses were performed as previously described (11, 12). For details, see Supplementary Information.

### Availability

Small RNA sequences and project have been deposited at the GEO under accession number GSE116122.

### ChIP and RT-qPCR experiments

Procedures, as a list of primer sequences used, were detailed in Supplementary Information.

## RESULTS

### Germline silencing induced at high temperature

Earlier studies of hybrid dysgenesis reported that high temperature enhances *P*-element repression (22) and that thermic modification of *P* repression can persist over several generations (23). Moreover, *P*-element repression in a strain carrying two *P*-elements inserted in a subtelomeric piRNA clusters can be stimulated by heat treatment (24). Very recently, the tracking of natural invasion of *P-element* in *Drosophila simulans* confirmed the key role of a high temperature in the establishment of repression through generations (25). These results suggested that temperature may influence piRNA cluster activity. In order to investigate whether high temperature (29°C) could affect the stability of *BX2* epialleles (*BX2^OFF^* and *BX2^ON^*) across generations, we generated flies carrying each of the *BX2* epialleles and a *P(TARGET)* reporter transgene sharing *P* and *lacZ* sequences with *BX2* on the same chromosome. As was previously described (11), at 25°C *BX2^OFF^* does not synthesized functional piRNAs complementary to *P(TARGET)* resulting in ß-Galactosidase expression in whole ovaries of *BX2^OFF^, P(TARGET)* lines (FIG 1A), whereas in *BX2^ON^*, *P(TARGET)* lines, functional *lacZ* piRNAs are synthesized in the germline where they specifically repress the *P(TARGET)* ß-Galactosidase expression (FIG 1B). Both *BX2^OFF^, P(TARGET)* and *BX2^ON^, P(TARGET)* lines incubated at 25°C for 23 generations maintained their epigenetic state showing that both epialleles are stable (FIG 1D, TABLE S1). At 29°C, the repression capacity of *BX2^ON^, P(TARGET)* lines remained stable through 25 generations whereas among the *BX2^OFF^, P(TARGET)* lines, 24.7% of females analyzed during 25 generations showed a complete and specific germline ß-Galactosidase repression (n=3812, FIG 1E and TABLE S1) suggesting a conversion of the *BX2^OFF^* epiallele into *BX2^ON^*. Interestingly, the appearance of females showing ß-Galactosidase repression was gradual and stochastic, resulting in a global frequency that increased with the number of generations (TABLE S1). To test whether the temperature-induced conversion was stable, a set of five lines showing full repression capacities after 23 generations at 29°C, obtained from an independent experiment, were transferred to 25°C and tested for their silencing capacities for several generations. In all cases, the silencing capacities of the *BX2^ON^* epiallele induced at 29°C remained stable during 50 additional generations at 25°C (TABLE S2). These stable *BX2^ON^* lines converted by high temperature were named hereafter *BX2^Θ^* (Greek theta for temperature) to distinguish them from the *BX2\** lines converted by maternally inherited piRNAs (11). Taken together, our data show that *BX2^OFF^* can be functionally converted by high temperature (29°C), strongly suggesting that *de novo* piRNA production can occur in the absence of maternal inheritance of homologous piRNAs.

**Figure 1.**
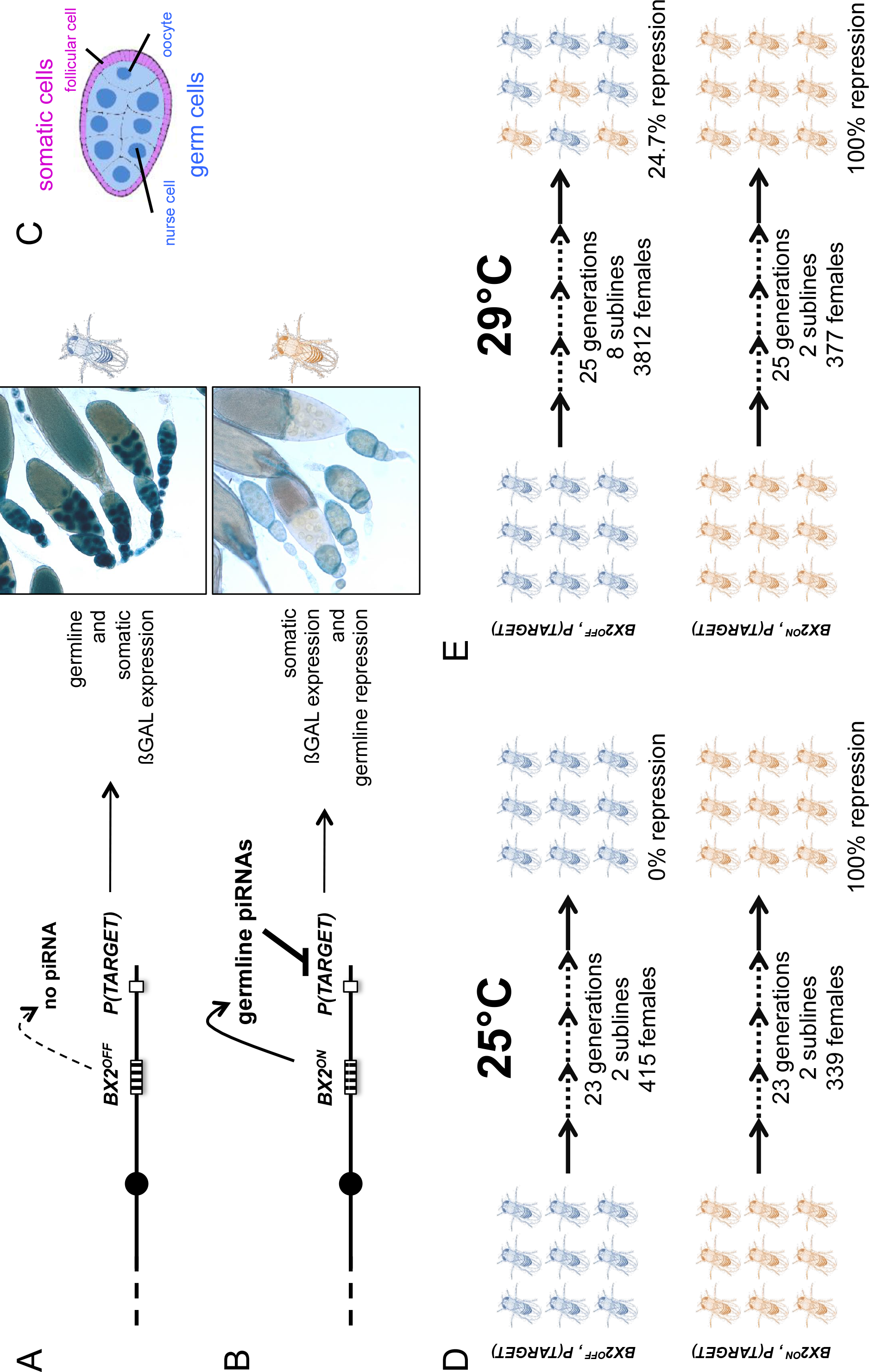
Functional assay of the *BX2* epigenetic state. Females carrying either one of the *BX2* epialleles and a *P(TARGET)* reporter transgene were analyzed. (A) When *BX2* is OFF for production of piRNA (*BX2^OFF^*), no repression of *P(TARGET)* occurs, allowing expression of ßGalactosidase in both germline and in somatic lineages in ovaries (ß-Galactosidase staining). *BX2^OFF^* is illustrated by a blue fly. (B) When *BX2* is ON for production of piRNA (*BX2^ON^*), repression of *P(TARGET)* occurs only in the germline lineage. *BX2^ON^* is illustrated by a light brown fly. (C) Drawing of an intermediate egg chamber showing germ cells (nurse cells and oocyte in blue) surrounded by somatic follicular cells (in pink), adapted from (43). (D) At 25°C, *BX2^OFF^* and *BX2^ON^* are stable over generations. (E) At 29°C, *BX2^OFF^* can be converted into *BX2^ON^*, while *BX2^ON^* is stable over generations.

### *BX2* lines converted by high temperature or by maternal homologous piRNA inheritance present identical functional and molecular properties

We further characterized the functional and molecular properties of *BX2^Θ^* activated by temperature and compared them to *BX2\** activated by maternal inheritance of homologous piRNAs. First, we compared the maternal and paternal *BX2* locus inheritance effect of three *BX2^Θ^* lines and three *BX2\** at 25°C (FIG S1A). Maternal inheritance of either *BX2^Θ^* or *BX2\** loci leads to complete and stable repression of ß-Galactosidase expression (n flies = 152 and 159, respectively, TABLE S3, whereas paternal inheritance of either *BX2^Θ^* or *BX2\** loci, *i.e*. in absence of maternal piRNA deposition, results in ß-Galactosidase expression, and thus a definitive loss of *BX2* silencing capacities (n flies = 156 and 155, respectively, TABLE S3). Second, we had shown before that the progeny having paternally inherited the *BX2^OFF^* locus and maternally inherited only the piRNAs produced by the *BX2\** mother without the genomic locus showed 100% conversion (11). This process of recurrent conversions of an allele that is heritable without DNA modification is known as paramutation, thus *BX2\** females are paramutagenic, *i.e*. able to trigger paramutation. To test this property on *BX2* lines converted by temperature, *BX2^Θ^* females were crossed with *BX2^OFF^* males. The progeny that inherited the paternal *BX2^OFF^* locus but not the maternal *BX2^Θ^* locus was selected and three independent lines were established (FIG S1B). Silencing measured during 20 generations revealed 100% of repression capacity showing that *BX2^Θ^* is also paramutagenic (n flies = 159, TABLE S4).

To determine whether the silencing capacities of *BX2^Θ^* involved piRNAs, small RNAs from *BX2^OFF^*, *BX2^Θ^* and *BX2\** ovaries were extracted and sequenced (TABLE S5). Unique reads matching the *P(lacW)* sequences were identified only in the *BX2^Θ^* and *BX2\** libraries (FIG 2A). Most of these small RNAs display all the characteristics of *bona fide* germline piRNAs, *i.e*. a high proportion of 23-29 nt with a strong U bias on the first 5’ nucleotide and an enrichment of 10-nucleotides overlap between sense and antisense piRNAs also known as the ping-pong signature (FIG 2B-C, (3, 4). As a control, the *42AB* piRNA cluster, a canonical germline dual-strand piRNA cluster, presented no significant difference between the three genotypes (FIG S2). Therefore, these results show that high temperature can initiate piRNA production from *BX2* naive sequences (*BX2^OFF^*) and strongly suggest that once a piRNA cluster is activated for piRNA production, the ‘ON’ state is maintained at each generation by maternal inheritance of piRNAs.

**Figure 2.**
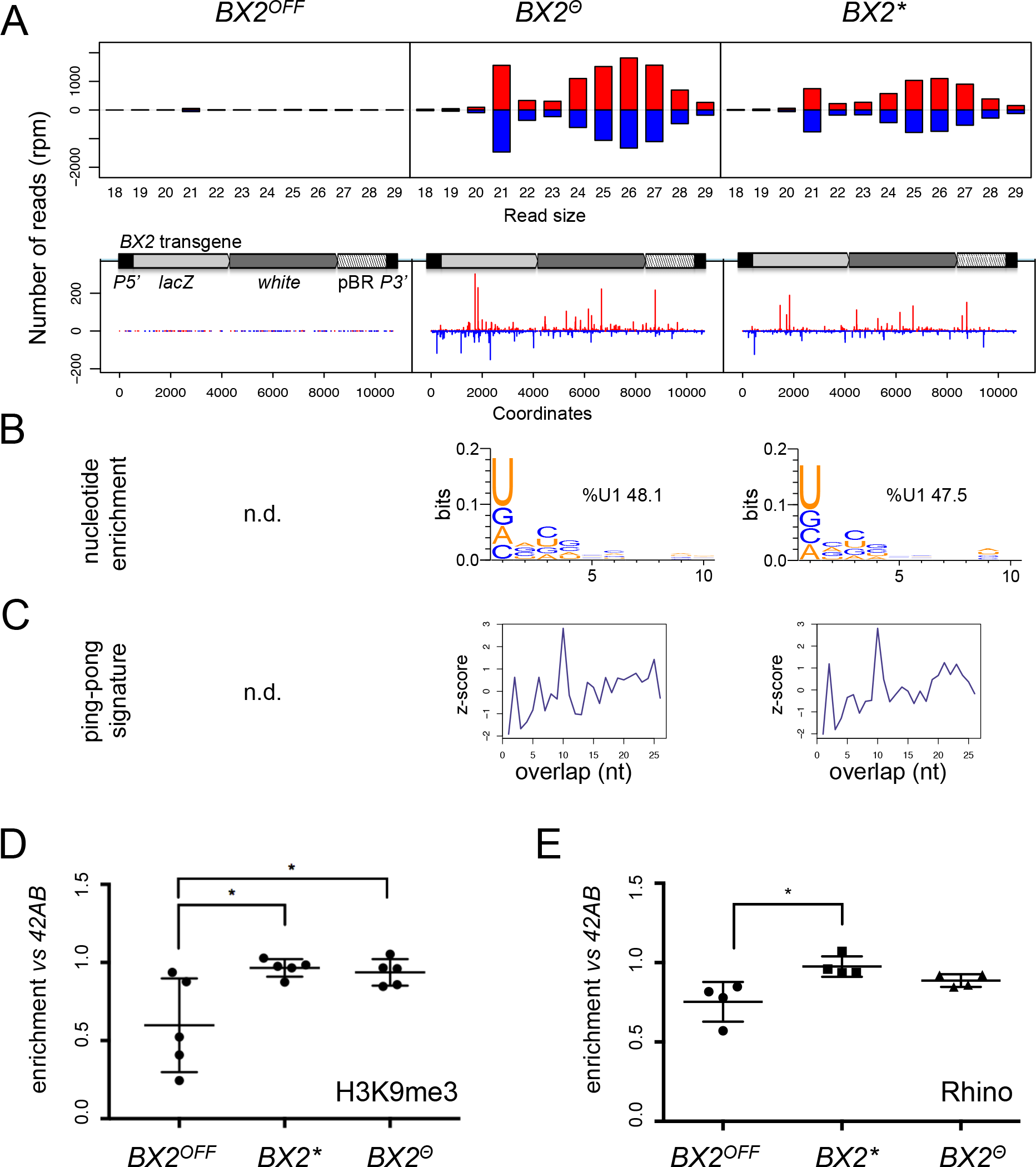
*BX2^Θ^* and *BX2\** produce piRNAs and are enriched in H3K9me3. (A) Size distribution of ovarian small RNA matching *BX2* transgene sequences reveals that both *BX2^Θ^* and *BX2\** but not *BX2^OFF^* produce 21-nt endo siRNAs and 23-29-nt piRNAs (upper panels, pBR = backbone plasmid pBR322). Positive and negative values correspond to sense (red) and antisense (blue) reads, respectively. Unique 23-29 nt mappers are shown on the *BX2* transgene sequences (lower panels). (B) Proportion of 23-29nt small RNAs from *BX2^Θ^* and *BX2\** matching transgene sequence with a U at the first position are shown. n.d.: not determined due to low number of reads. (C) Relative frequency (z-score) of overlapping sense-antisense 23-29nt RNA pairs reveals an enrichment of 10-nucleotides overlapping corresponding to the ping-pong signature. (D) H3K9me3 and (E) Rhino binding on the *BX2* transgene in ovaries of *BX2^OFF^*, *BX2^Θ^* and *BX2\** strains revealed by chromatin immunoprecipitation (ChIP) followed by quantitative PCR (qPCR) on specific *white* sequences. In both "ON" strains, *BX2^Θ^* and *BX2\**, H3K9me3 and Rhino levels over the transgene are very similar and higher than in the *BX2^OFF^* strain (* = P < 0.05, Mann– Whitney–Wilcoxon test, n = 5).

To test whether other non-piRNA producing genomic loci have started to produce piRNAs following high temperature treatment, we looked for specific piRNAs (23-29 nt) matching at unique positions on *Drosophila* chromosomes and compared them between *BX2^Θ^* and *BX2^OFF^*. The reads were then resampled per 50 kilobases windows. To eliminate background noise, only regions that produced more than 5 piRNAs per kilobase on average on both libraries were considered. Only exons of the *white* gene present in the *P(lacW)* transgenes of *BX2* showed differential piRNA expression (log2 ratio > 8.5, FIG S3). This analysis revealed that the activation of piRNA production after thermic treatment is restricted to the *BX2* locus, suggesting that all other loci able to produce piRNAs are already activated.

Previous studies had suggested that the chromatin state plays a role in the differential activity of *BX2* (26). We therefore profiled H3K9me3 marks and Rhino binding on the *P(lacW)* transgene in ovaries from *BX2^OFF^*, *BX2^Θ^* and *BX2\** strains by chromatin immunoprecipitation (ChIP) followed by quantitative PCR (qPCR). In both strains *BX2^Θ^* and *BX2\**, H3K9me3 and Rhino were similarly enriched over the *P(lacW)* transgene compared to the *BX2^OFF^* strain (significantly for H3K9me3, FIG 2D, 2E). Taken together, all these results show that *de novo* activation of *BX2^OFF^* by 29°C treatment (*BX2^Θ^*) or paramutation by maternal inheritance of homologous piRNAs (*BX2\**) lines leads to similar functional and molecular properties.

### Epigenetic conversion at 29°C occurs at a low rate from the first generation

To explain the low occurrence and the generational delay of *BX2* conversion at 29°C (TABLE S1) we propose that conversion is a complete but rare event occurring in a small number of egg chambers at each generation. Under this hypothesis, the sampling size of tested females should be crucial to observe such stochastic events. We therefore increased the number of analyzed females raised at 29°C during one generation. For this, eggs laid by females maintained at 25°C carrying the *P(TARGET)* reporter transgene and the *BX2^OFF^* locus were collected during three days. These eggs were then transferred at 29°C until adults emerged (FIG 3A). In order to follow their offspring, we crossed individually 181 G1 females with two siblings and let them lay eggs for three days at 25°C. These 181 G1 females were then stained for ß-Galactosidase expression. Strikingly, repression occurred only after one generation at 29°C in a few of the egg chambers of 130 G1 females ovaries (≈2.7% of the estimated total number of G1 egg chambers n≈21700, FIG 3B, right panel). These results favor our hypothesis where epigenetic conversion of *BX2^OFF^* into *BX2^Θ^* is an instantaneous and complete event occurring at low frequency per egg chamber and at each generation that is kept at 29°C.

**Figure 3.**
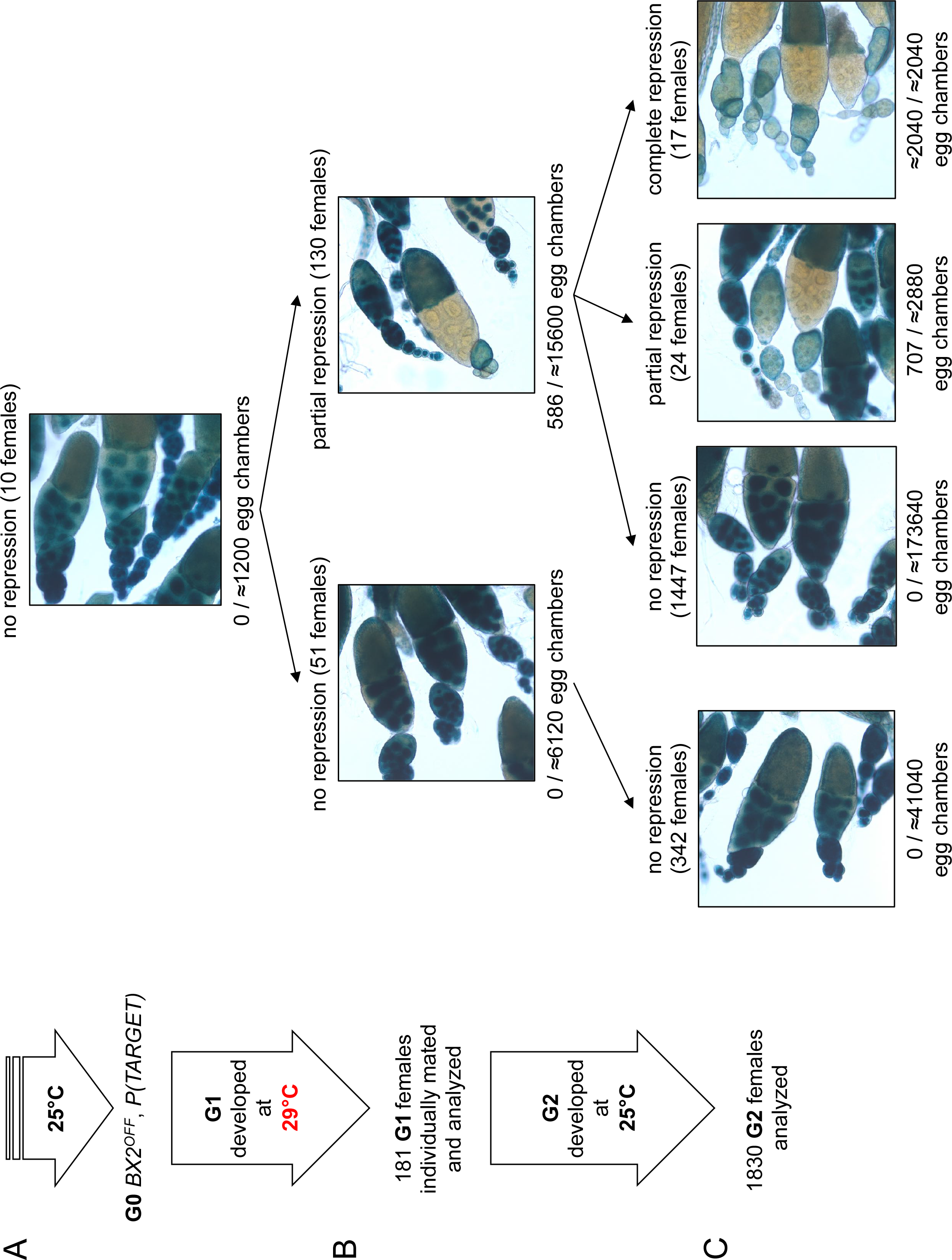
*BX2* conversion at 29°C occurs in one generation at a low rate. (A) G0 females carrying the *P(TARGET)* reporter and *BX2^OFF^* laid eggs at 25°C during three days. The *BX2^OFF^* state of these females was confirmed after the three days at 25°C by ß-Galactosidase staining (number of egg chambers ≥ 1200). (B) Their eggs were allowed to develop at 29°C until emergence of the next generation. G1 females (n = 181) were individually mated with two siblings and left to lay for three days at 25°C. G1 females were then individually stained for ßGalactosidase expression. Strikingly, 130 females (71.8%) show ß-Galactosidase repression in some egg chambers (586 among ≈21700 - estimation of the total egg chamber number among 181 females). The *BX2^OFF^* into *BX2^ON^* conversion frequency is ≈2.7%. (C) Analysis of each G1 female progeny developed at 25°C by ß-Galactosidase staining. The progeny of the 51 G1 females that did not present repression maintained *BX2^OFF^* state (n flies = 342). Most of the progeny of the 130 G1 females presenting conversion show no repression (97.2%, n flies = 1488) while 41 females present partial (n = 24) or complete (n = 17) repression of the germline expression of ß-Galactosidase.

To test the stability of the epigenetic *BX2^Θ^* states observed in G1 females, offspring daughters (G2) were raised at 25°C and their ovaries examined for ß-Galactosidase expression. Among G2 females, partial (n=24) or complete (n=17) repression of ß-Galactosidase expression in the germline was observed only in the progeny of those 130 G1 females in which partial repression was previously detected (FIG 3C). The proportion of 2.2% of converted G2 females (41/1830) is reminiscent with the proportion of repressed egg chambers observed in G1. The progeny of the 51 G1 females that did not present repression (FIG 3B left panel) did not show spontaneous conversion (FIG 3C). Taken together, these observations strongly suggest that newly converted *BX2^ON^* egg chambers give rise to adult females with complete or partial silencing capacities. The low conversion rate observed in thousands of flies after one generation raised at high temperature and its stability through the next generation might explain the apparent delay of *BX2^ON^* conversion of dozens of flies continuously raised at 29°C observed in the first set of experiment (see FIG 1E and TABLE S1). This hypothesis is in agreement with a theoretical model that we tested on eight independent lines followed during 25 generations at 29°C (TABLE S6, FIG S4 and SUPPLEMENTAL INFORMATION 1).

Altogether, our results illustrate how environmental modifications like high temperature experienced during one generation might stably modify the epigenome of the future ones. Such newly acquired epigenetic state may spread in a given population within a few generations.

### High temperature increases *BX2* antisense RNA but not piRNAs

Previously, we showed that *BX2^OFF^* and *BX2^ON^* produce similar amounts of sense and antisense transcripts (11). However, these transcripts do not lead to ß-Galactosidase expression in the germline nor piRNA production in the *BX2^OFF^* line. We wondered if high temperature might change the RNA steady-state level of *BX2*. To ensure that we were detecting RNA specifically from *BX2*, qRT-PCR experiments targeting *lacZ* gene were carried out on ovarian RNA extracted from *BX2^OFF^* line that did not contain *P(TARGET)*. We observed a significant increase in the steady-state *BX2* RNA levels at 29°C compared to 25°C (FIG 4A). Remarkably, strand-specific qRT-PCR experiments revealed that only *BX2* antisense transcripts increase at 29°C (FIG 4B). As *BX2* is inserted into the *AGO1* gene in a convergent transcription manner (11), FIG 4C), we compared *AGO1* steady-state RNA level at 29°C and 25°C. *AGO1* presents a significant increase at 29°C (FIG 4D) suggesting that an increase of transcription from the *AGO1* promoter at 29°C could lead to an increase of *BX2* antisense RNA transcription.

**Figure 4.**
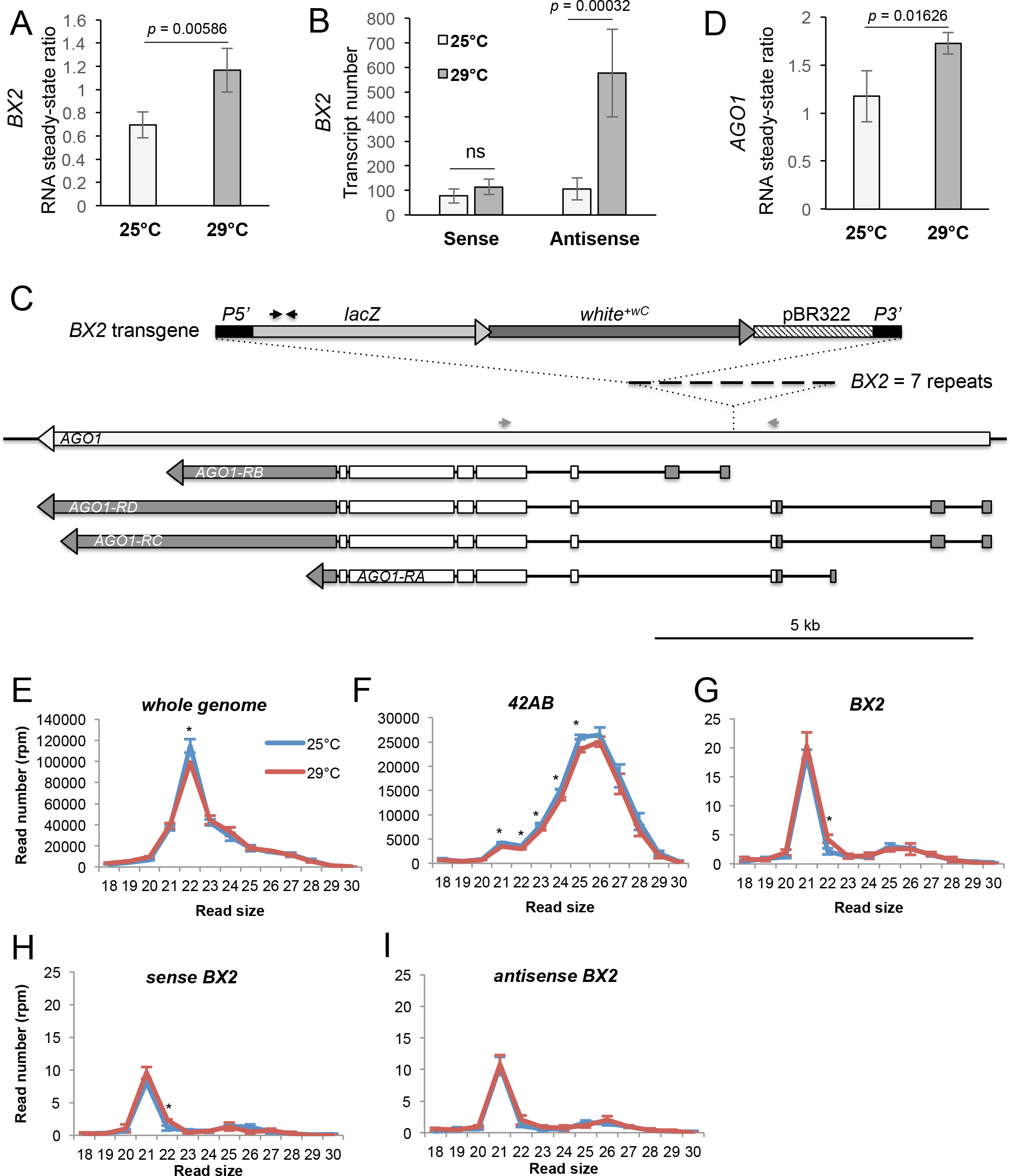
*BX2^OFF^* antisense RNA increase at 29°C. (A) RT-qPCR experiments revealed that the steady-state level of ovarian *lacZ* RNA from *BX2* are more abundant at 29°C (n=5) compared to 25°C (n=6). (B) Sense-specific RT-qPCR experiments revealed that only antisense transcripts from *BX2* are increased (25°C n=6, 29°C n=4). Significant *p*-values are given (bilateral Student’s *t*-test). ns: not significant. (C) Map of the *BX2* locus containing 7 *P(lacW)* transgenes inserted into the *AGO1* gene. *P(lacW)* and *AGO1* are drawn to scale. The *lacZ* gene contained in *P(lacW)* and *AGO1* are transcriptionally in opposite direction. Black arrows show *lacZ* primers used for (A) and (B) experiments. Grey arrows show *AGO1* primers used for (D) experiment. (D) RT-qPCR experiments performed on flies devoid of *P(lacW)* transgenes revealed that the steady-state level of ovarian RNA from *AGO1* are more abundant at 29°C (n=5) compared to 25°C (n=6). (EI) To compare small RNAs at 25 *versus* 29°C, total RNA were extracted from *BX2^OFF^* ovaries dissected from adults incubated at 25°C or 29°C. Three samples were tested for each temperature. Small RNAs from 18 to 30 nucleotides were deep sequenced. For each library, normalization has been performed for 1 million reads matching *Drosophila* genome (rpm, TABLE S7). Size distributions of unique reads that match reference sequences are given. (E) Small RNAs matching *Drosophila* genome present similar profiles in both temperatures except for 22nt RNA that are more represented at 25°C. (F) The 21 to 25nt reads matching the *42AB* piRNA cluster that range from 21 to 25nt are slightly more abundant at 25°C. (G) Strikingly, almost only 21nt RNAs match *BX2* sequence. They are equally distributed among sense (H) and antisense (I) sequences at both temperatures. * = p < 0.05, bilateral Student’s *t*-test.

We then examined whether the increase of *BX2* antisense RNAs leads to an increase of antisense small RNAs. Ovarian small RNAs (18 to 30 nucleotides) of three biological replicates of *BX2^OFF^* flies (without *P(TARGET)*) raised at 25°C and at 29°C for one generation were sequenced and the read numbers normalized (TABLE S7). A slight significant decrease could be observed at 29°C for small RNAs matching the whole genome (FIG 4E) or the *42AB* piRNA cluster (FIG 4F). Strikingly, no piRNAs were produced from the *BX2* locus at 25°C nor after one generation at 29°C. Thus, the increase of *BX2* antisense transcripts observed at 29°C (FIG 4B) did not correlate with an increase of corresponding antisense piRNAs. At 25°C and 29°C, *BX2^OFF^* produced the same little amount of 21nt small RNAs, equally distributed between sense and antisense (FIG 4G, H, I) suggesting that *BX2* transcripts are processed into siRNAs.

### Heat conversion requires a homologous sequence *in trans*

qRT-PCR experiments described above were carried out on flies bearing only the *BX2* locus while all conversion experiments at 29°C were done with flies bearing the *BX2* locus and the *P(TARGET)*. We next asked whether the *P(TARGET)* transgene could participate in the conversion process of *BX2*. For this, ‘heat-activated-conversion’ experiments of *BX2* were done in flies not carrying the *P(TARGET)* (FIG S5). To assess the *BX2* epigenetic state of the G1 raised at 29°C, 157 G1 females were then individually crossed at 25°C with males harboring the *P(TARGET)* transgene. Among the 1137 G2 females analyzed, only one female presented partial repression of the ß-Galactosidase expression and none presented complete repression (FIG S5). If we compare these results with those obtained with the *BX2*, *P(TARGET)* lines (FIG 3), the difference was highly significant (*p* = 8.5.10^−6^, homogeneity chi2 test, TABLE 1). To further validate the requirement of the euchromatic homologous transgene *P(TARGET)* in establishing the temperature-dependent *BX2* conversion, we generated eight independent lines in which *BX2* was recombined into the same *P(TARGET)* genetic background but without the *P(TARGET)* transgene. After 13 generations at 29°C, no female showing ß-Galactosidase repression was observed (TABLE S8), arguing against a background effect of the *P(TARGET)* line in the conversion phenomenon (*p*=7.04.10^−^44, homogeneity chi2 test compared to *BX2, P(TARGET)* after 13 generations at 29°C, TABLE S1). We concluded that the presence of a reporter transgene sharing homologous sequences (*i.e*. *lacZ*) with *BX2* is essential for the *BX2* conversion at 29°C.

**Table 1.** *P(TARGET)* requirement in the *BX2* conversion process.

| Repression occurrence | Number of G2 females |  |  |
| --- | --- | --- | --- |
|  | none | partial | complete |
| *with P(TARGET)* | 1789 | 24 | 17 |
| *without P(TARGET)* | 1136 | 1 | 0 |

To summarize, in the absence of *P(TARGET)* sequences, *BX2^OFF^* produces more antisense transcripts, no piRNAs and is unable to be converted to *BX2^ON^* at 29°C. In contrast, when *P(TARGET)* is present, *BX2* conversion and piRNA production are observed at 29°C. We propose that, at 29°C, the interaction between the excess of *BX2* antisense transcripts and the sense *P(TARGET)* transcripts is a prerequisite for the production of *de novo* piRNAs and the conversion of *BX2* into an active piRNA cluster.

## DISCUSSION

Here, we report on the heritable establishment of a new piRNA cluster associated with silencing properties induced by high temperature during development. By doing so, we uncovered that thermic changes, like global warming, might have consequences on the efficiency of TE regulation. The epigenetic response to heat exposure has been studied in several model species: in *Arabidopsis* for instance, increasing temperature induces transcriptional activation of repetitive elements (27–29). Whether these changes involve chromatin modifications is not clear but none of these modifications have been found to be heritable through generations except in mutants for siRNA biogenesis where high frequency of new TE insertions was observed in the progeny of stressed plants (27). In animals, response to heat can result in modification of DNA methylation at specific loci in reef building coral (30), chicken (31) and wild guinea pigs (32). In the latter, modifications affecting ≈50 genes are inherited in G1 progeny (32). However, the mechanisms of this heritability are not yet understood. In *Drosophila*, heat-shock treatment of 0-2h embryos for one hour at 37°C or subjecting flies to osmotic stress induce phosphorylation of dATF-2 and its release from heterochromatin (33). This defective chromatin state is maintained for several generations before returning to the original state. We have tested if such stresses were able to convert *BX2^OFF^* into *BX2^ON^* in one generation but neither heat-shock nor osmotic stress induces *BX2* conversion (TABLE S9), suggesting that *BX2* activation does not depend on *dATF-2*. In a more recent paper, Fast *et al*. (34) found less piRNAs at 29°C than at 18°C. However, RNAseq analyses of differentially expressed genes involved in the piRNA pathway were not conclusive, as some piRNA genes were more expressed at 29°C (*ago3*, *aub*, *zuc, armi*) while others were less expressed (*shu*, *hsp83, Yb*) (34). Overall, the enhancement of the piRNA ping-pong amplification loop observed at 29°C was attributed to RNA secondary structures, because of a lack of specificity for any particular class of TE (34). Furthermore, in contrast to our RT-qPCR results obtained at 25°C and 29°C (FIG 4D), *AGO1* was not differentially expressed between 18°C and 29°C (34). This difference can be explained, because, in our experiments, we checked for specific spliced transcripts of *AGO1* that originate upstream the *BX2* insertion point (FIG 4C). Taken together, all these observations show that temperature modification might induce epigenetic changes in several species but the underlying mechanisms remain largely unsolved.

Whole genome comparison of small RNA sequencing between *BX2^OFF^* and newly *BX2^ON^* heat-converted flies (*BX2^Θ^*) did not reveal additional regions stably converted for piRNA production (FIG S3). This suggests that no other loci are metastable for piRNA production, *i.e*. all potential piRNA clusters are already activated at 25°C. This observation raises the question of what makes *BX2* locus competent for piRNA activation at high temperature. The *BX2* locus is the result of successive induced transpositions of *P(lacW)* used to screen for *white*-variegating phenotype (20). Thus, *BX2* resembles natural TE clusters where TEs have the capacity to transpose into each other assembling structures named *nested TE*, as described in numerous genomes (35–37). Some *nested TE* loci might have the capacity to be activated and respond to a new TE invasion. In *Drosophila*, *BX2*-like tandemly inserted transgenes were shown to be new sites of HP1 enrichment in larval salivary glands emphasizing a heterochromatic structure at the *BX2* locus in somatic cells (38) and to cause pairing-dependent silencing (39). However, when tested for repressing capacities, this strain is inactive for *BX2* piRNA production (11, 40). We have shown that maternally inherited *P(lacW)* piRNAs are able to paramutate with complete and stable penetrance from an inactive *BX2* locus into an active locus for piRNA production (11). The paramutated *BX2* locus appeared to be a genuine piRNA cluster since it is sensitive to a number of factors known to be involved in piRNA biogenesis such as *aub*, *rhi*, *cuff*, *zuc* (12) and *moonshiner* (FIG S6). The number of transgene copies appears to be crucial in the processes since smaller number of transgenes results in lesser somatic heterochromatinization (20, 38), lesser pairing-dependent silencing (39) and unstable paramutation (11). Taken together, these data suggest that the heterochromatic structure of a cluster precedes piRNA production. This is sustained by our ChIP experiments showing that H3K9me3 levels on *BX2^OFF^* are slightly below the H3K9me3 level of piRNA producing states (*BX2^ON^* and *BX2^Θ^* FIG 2D). The same observation can be made for Rhino (FIG 2E) suggesting that Rhino may be already present on the *BX2^OFF^* locus but below the threshold required for piRNA production as suggested by (41). Thus, a locus made of repeated sequences and being likely heterochromatic (H3K9me3, Rhino) is a necessary but not sufficient condition to specify an active piRNA cluster.

In the germline, piRNA clusters produce piRNAs from both strands and it was recently shown that, in most cases, transcription initiates within clusters on both strands through the interaction of Rhino and Moonshiner (9). In few cases, however, piRNA cluster transcription may take advantage of the read-through from a flanking promoter (9). Zhang *et al*. (8) have shown that tethering Rhino onto a transgene leads to its repression but the production of piRNA depends on the presence of another transgene producing antisense RNA. Moreover, in the context of the *Pld* promoter deletion, a gene flanking the *42AB* piRNA cluster, flies can produce *Pld* piRNAs only if a *Pld* cDNA is expressed *in trans* (9). From all these observations, it emerges that production of simultaneous sense and antisense RNA is a shared requirement for piRNA production. However, even if *BX2^OFF^* is transcribed on both strands, it still remains inactive for piRNA production.

In addition to having a number of heterochromatic repeats and a double stranded transcription, the production of *de novo* piRNAs from *BX2* requires a triggering signal. From our experiments, *BX2* conversion relies on the simultaneous increase of both sense and antisense RNAs. An active role of euchromatic copies in the establishment of new piRNA clusters by high temperature appears to be consistent with what would naturally happen during the invasion of a naive genome by new TEs or when chromosomal breakages occur leading to the loss of piRNA cluster loci (42). At first, uncontrolled euchromatic TE transposition takes place before the establishment of repression. Such repression would occur after a copy integrates into a preexisting piRNA cluster or by the generation of a new cluster made by successive nested copies. Consequently, clusters of elements cannot exist without transcriptionally active euchromatic copies. The increase of germline antisense transcripts upon stress or environmental factors, depending on the neighboring genomic environment, and the concomitant presence of numerous sense transcripts from euchromatic active copies appear to be the starting signal for new piRNA production. These piRNAs can then be inherited at the next generation where they will stably paramutate the corresponding DNA locus with repetitive nature. At that time, the triggering signal is no longer necessary since *BX2* remains activated once flies get back at 25°C. Future generations thus remember what was once considered a threat only through the legacy of maternal piRNAs.

## ACKNOWLEDGEMENTS

We thank Doug Dorer, Steve Henikoff, Julius Brennecke and the *Bloomington Stock Center* for providing stocks. We thank Bill Theurkauf for providing antibodies. We thank flybase.org for providing databases. We thank Ritha Zamy for technical assistance. We thank Clément Carré, Ana Maria Vallès and Jean-René Huynh for critical reading of the manuscript. We thank Christophe Antoniewski for helpful advices and development of the ARTbio platform (http://artbio.fr/).

## REFERENCES

1. Wylie A, Jones AE, & Abrams JM (2016) p53 in the game of transposons. Bioessays 38(11):1111–1116.

2. Wylie A, et al. (2016) p53 genes function to restrain mobile elements. Genes Dev 30(1):64–77.

3. Brennecke J, et al. (2007) Discrete small RNA-generating loci as master regulators of transposon activity in Drosophila. Cell 128(6):1089–1103.

4. Gunawardane LS, et al. (2007) A slicer-mediated mechanism for repeat-associated siRNA 5’ end formation in Drosophila. Science 315(5818):1587–1590.

5. Le Thomas A, et al. (2013) Piwi induces piRNA-guided transcriptional silencing and establishment of a repressive chromatin state. Genes Dev 27(4):390–399.

6. Sienski G, Donertas D, & Brennecke J (2012) Transcriptional silencing of transposons by Piwi and maelstrom and its impact on chromatin state and gene expression. Cell 151(5):964–980.

7. Mohn F, Sienski G, Handler D, & Brennecke J (2014) The rhino-deadlock-cutoff complex licenses noncanonical transcription of dual-strand piRNA clusters in Drosophila. Cell 157(6):1364–1379.

8. Zhang Z, et al. (2014) The HP1 homolog rhino anchors a nuclear complex that suppresses piRNA precursor splicing. Cell 157(6):1353–1363.

9. Andersen PR, Tirian L, Vunjak M, & Brennecke J (2017) A heterochromatin-dependent transcription machinery drives piRNA expression. Nature 549(7670):54–59.

10. Brennecke J, et al. (2008) An epigenetic role for maternally inherited piRNAs in transposon silencing. Science 322(5906):1387–1392.

11. de Vanssay A, et al. (2012) Paramutation in Drosophila linked to emergence of a piRNA-producing locus. Nature 490(7418):112–115.

12. Hermant C, et al. (2015) Paramutation in Drosophila Requires Both Nuclear and Cytoplasmic Actors of the piRNA Pathway and Induces Cis-spreading of piRNA Production. Genetics 201(4):1381–1396.

13. Marie PP, Ronsseray S, & Boivin A (2017) From Embryo to Adult: piRNA-Mediated Silencing throughout Germline Development in Drosophila. G3 (Bethesda) 7(2):505–516.

14. Muerdter F, et al. (2012) Production of artificial piRNAs in flies and mice. RNA 18(1):42–52.

15. Poyhonen M, et al. (2012) Homology-dependent silencing by an exogenous sequence in the Drosophila germline. G3 (Bethesda) 2(3):331–338.

16. Klattenhoff C, et al. (2009) The Drosophila HP1 homolog Rhino is required for transposon silencing and piRNA production by dual-strand clusters. Cell 138(6):1137–1149.

17. Huang X, Fejes Toth K, & Aravin AA (2017) piRNA Biogenesis in Drosophila melanogaster. Trends Genet 33(11):882–894.

18. Brink RA (1956) A Genetic Change Associated with the R Locus in Maize Which Is Directed and Potentially Reversible. Genetics 41(6):872–889.

19. Chandler VL (2007) Paramutation: from maize to mice. Cell 128(4):641–645.

20. Dorer DR & Henikoff S (1994) Expansions of transgene repeats cause heterochromatin formation and gene silencing in Drosophila. Cell 77(7):993–1002.

21. Lemaitre B, Ronsseray S, & Coen D (1993) Maternal repression of the P element promoter in the germline of Drosophila melanogaster: a model for the P cytotype. Genetics 135(1):149–160.

22. Ronsseray S, Anxolabehere D, & Periquet G (1984) Hybrid dysgenesis in Drosophila melanogaster: influence of temperature on cytotype determination in the P-M system. Mol Gen Genet 196(1):17–23.

23. Ronsseray S (1986) P-M System of Hybrid Dysgenesis in Drosophila-Melanogaster - Thermal Modifications of the Cytotype Can Be Detected for Several Generations. Molecular & General Genetics 205(1):23–27.

24. Ronsseray S, Lehmann M, & Anxolabehere D (1991) The maternally inherited regulation of P elements in Drosophila melanogaster can be elicited by two P copies at cytological site 1A on the X chromosome. Genetics 129(2):501–512.

25. Kofler R, Senti KA, Nolte V, Tobler R, & Schlotterer C (2018) Molecular dissection of a natural transposable element invasion. Genome Res 28(6):824–835.

26. Le Thomas A, et al. (2014) Transgenerationally inherited piRNAs trigger piRNA biogenesis by changing the chromatin of piRNA clusters and inducing precursor processing. Genes Dev 28(15):1667–1680.

27. Ito H, et al. (2011) An siRNA pathway prevents transgenerational retrotransposition in plants subjected to stress. Nature 472(7341):115–119.

28. Pecinka A, et al. (2010) Epigenetic regulation of repetitive elements is attenuated by prolonged heat stress in Arabidopsis. Plant Cell 22(9):3118–3129.

29. Tittel-Elmer M, et al. (2010) Stress-induced activation of heterochromatic transcription. PLoS Genet 6(10):e1001175.

30. Dimond JL & Roberts SB (2016) Germline DNA methylation in reef corals: patterns and potential roles in response to environmental change. Mol Ecol 25(8):1895–1904.

31. Yossifoff M, Kisliouk T, & Meiri N (2008) Dynamic changes in DNA methylation during thermal control establishment affect CREB binding to the brain-derived neurotrophic factor promoter. Eur J Neurosci 28(11):2267–2277.

32. Weyrich A, et al. (2016) Paternal intergenerational epigenetic response to heat exposure in male Wild guinea pigs. Mol Ecol 25(8):1729–1740.

33. Seong KH, Li D, Shimizu H, Nakamura R, & Ishii S (2011) Inheritance of stress-induced, ATF-2-dependent epigenetic change. Cell 145(7):1049–1061.

34. Fast I, et al. (2017) Temperature-responsive miRNAs in Drosophila orchestrate adaptation to different ambient temperatures. RNA 23(9):1352–1364.

35. Gao C, et al. (2012) Characterization and functional annotation of nested transposable elements in eukaryotic genomes. Genomics 100(4):222–230.

36. Liu R, et al. (2007) A GeneTrek analysis of the maize genome. Proc Natl Acad Sci U S A 104(28):11844–11849.

37. Zanni V, et al. (2013) Distribution, evolution, and diversity of retrotransposons at the flamenco locus reflect the regulatory properties of piRNA clusters. Proc Natl Acad Sci U S A 110(49):19842–19847.

38. Fanti L, Dorer DR, Berloco M, Henikoff S, & Pimpinelli S (1998) Heterochromatin protein 1 binds transgene arrays. Chromosoma 107(5):286–292.

39. Dorer DR & Henikoff S (1997) Transgene repeat arrays interact with distant heterochromatin and cause silencing in cis and trans. Genetics 147(3):1181–1190.

40. Josse T, et al. (2008) Telomeric trans-silencing in Drosophila melanogaster: tissue specificity, development and functional interactions between non-homologous telomeres. PloS one 3(9):e3249.

41. Akulenko N, et al. (2018) Transcriptional and chromatin changes accompanying de novo formation of transgenic piRNA clusters. RNA 24(4):574–584.

42. Asif-Laidin A, et al. (2017) Short and long-term evolutionary dynamics of subtelomeric piRNA clusters in Drosophila. DNA Res 24(5):459–472.

43. Frydman HM & Spradling AC (2001) The receptor-like tyrosine phosphatase lar is required for epithelial planar polarity and for axis determination within drosophila ovarian follicles. Development 128(16):3209–3220.

